# Gene expression analyses reveal differences in children’s response to malaria according to their age

**DOI:** 10.1101/2023.10.24.563751

**Authors:** Kieran Tebben, Salif Yirampo, Drissa Coulibaly, Abdoulaye K. Koné, Matthew B. Laurens, Emily M. Stucke, Ahmadou Dembélé, Youssouf Tolo, Karim Traoré, Amadou Niangaly, Andrea A. Berry, Bourema Kouriba, Christopher V. Plowe, Ogobara K Doumbo, Kirsten E. Lyke, Shannon Takala-Harrison, Mahamadou A. Thera, Mark A. Travassos, David Serre

## Abstract

In Bandiagara, Mali, children experience on average two clinical malaria episodes per season. However, even in the same transmission area, the number of uncomplicated symptomatic infections, and their parasitemia, vary dramatically among children. To examine the factors contributing to these variations, we simultaneously characterized the host and parasite gene expression profiles from 136 children with symptomatic falciparum malaria and analyzed the expression of 9,205 human and 2,484 *Plasmodium* genes. We used gene expression deconvolution to estimate the relative proportion of immune cells and parasite stages in each sample and to adjust the differential gene expression analyses. Parasitemia explained much of the variation in both host and parasite gene expression and revealed that infections with higher parasitemia had more neutrophils and fewer T cells, suggesting parasitemia-dependent neutrophil recruitment and/or T cell extravasation to secondary lymphoid organs. The child’s age was also strongly correlated with gene expression variations. *Plasmodium falciparum* genes associated with age suggested that older children carried more male gametocytes, while host genes associated with age indicated a stronger innate response (through TLR and NLR signaling) in younger children and stronger adaptive immunity (through TCR and BCR signaling) in older children. These analyses highlight the variability in host responses and parasite regulation during *P. falciparum* symptomatic infections and emphasize the importance of considering the children’s age when studying and treating malaria infections.

**One Sentence Summary:** Human and *P. falciparum* gene expression differs according to the infection’s parasitemia and the child’s age, highlighting an age-dependent response to malaria and complex cellular and molecular -host/parasite interactions.

## INTRODUCTION

Despite decades of progress towards elimination, malaria remains a major public health problem in endemic areas (*1*). In 2021, there were 247 million cases of malaria worldwide, resulting in over 600,000 deaths, mostly in children younger than five years old (*1*). Malaria is caused by *Plasmodium* parasites that are transmitted by *Anopheles* mosquitos (*2*). While five *Plasmodium* species cause human malaria – *P. falciparum, P. vivax, P. ovale, P. malariae* and *P. knowlesi* – *P. falciparum* is responsible for most cases of malaria worldwide (*2*) and causes the most severe forms of the disease (*3*). All symptoms of malaria stem from asexual replication of parasites in the blood, and therefore gene expression analysis of blood samples from infected patients can provide invaluable insights into the role of different host and parasite factors in regulating the disease. Several studies have previously used this approach to study human immune cells, revealing modulation of gene pathways regulated by pro-inflammatory cytokines over repeated malaria exposures in Malian adults (*4*), and development of memory B cells with atypical gene expression patterns over repeated exposures (*5*). Additionally, gene expression analysis has been used to study *Plasmodium* parasites, revealing parasitemia-dependent regulation of metabolism and cell death genes (*6*) and coordination of *var* gene expression (*7*). Few studies however have examined host and parasite transcripts from the same samples, and have done so specifically in the context of malaria disease severity: one study identified unique host and parasite gene expression patterns according to specific severe malaria complications (e.g., coma, hyperlactemia) and noted that human gene expression during severe malaria was driven by parasite load (*8*), while another identified activation of the innate immune system according to disease severity (*9*). However, it is important to understand how gene expression varies even in the context of uncomplicated disease.

One ubiquitous limitation of gene expression analyses of whole blood samples derives from the cellular heterogeneity of the material studied. Specifically, it is often difficult to determine if apparent gene expression differences among samples are i) genuine differences caused by differential gene regulation or, ii) caused by variations, among patients, in the relative proportions of white blood cell subsets or parasite developmental stages. Single cell RNA-sequencing or flow cytometry prior to bulk RNA-seq can circumvent this issue, but these methods are difficult to implement in field settings and can be prohibitively expensive in studying large cohorts. A commonly used approach to circumvent this issue, for parasite gene expression analysis, is to use synchronized *in vitro* cultures. However, the resulting gene expression profiles may be affected by the culture conditions and the absence of immune pressure and may therefore not accurately replicate the patterns observed *in vivo* (*10*) (*11*). Alternatively, one can computationally infer the proportions of different cells in a sample directly from the RNA-seq data using gene expression deconvolution (*12*): by comparing the normalized gene expression of a bulk RNA-seq experiment with the reference gene expression profiles of known cells, one can robustly estimate the proportions of both human immune cells (*12–14*), and *Plasmodium* developmental stages (*15–17*) present in a given whole blood sample.

Here, we used dual RNA sequencing (RNA-seq) to simultaneously characterize host and parasite gene expression from 136 whole-blood samples from Malian children during a symptomatic, uncomplicated *P. falciparum* infection. After estimating the relative proportions of the different immune cells and parasite developmental stages in each sample by gene expression deconvolution, we evaluated the contributions of different clinical and demographic parameters to the host and parasite gene expression profiles and determined whether gene expression differences were caused by true differences in gene expression or by changes in cell composition.

## RESULTS AND DISCUSSION

### Comprehensive profiling of host and parasite transcriptomes by dual RNA sequencing

We analyzed 136 whole-blood samples collected from children ages 1 to 15 years enrolled in a longitudinal incidence study in Bandiagara, Mali from 2009 – 2013 (*18*). All samples included were collected during an uncomplicated symptomatic *P. falciparum* malaria episode, defined by the study physicians as an unscheduled visit initiated by the patient in response to malaria symptoms (fever, headaches, joint pain, vomiting, diarrhea, or abdominal pain) with microscopic evidence of parasites (*18*). Children of both sexes were included in roughly equal proportions and most participants were of Dogon ethnicity (**Table 1**). For a subset of 120 individuals with at least two years of follow-up, we were able to determine i) the normalized number of subsequent malaria episodes and ii) the time to the next malaria episodes (weighted by the monthly risk of malaria and the child’s total time in the study, see **Materials and Methods** for details). Values for all variables considered in this study, as well as the date of sample collection for each sample, are available in **Supplemental Table 1.**

**Table 1.**
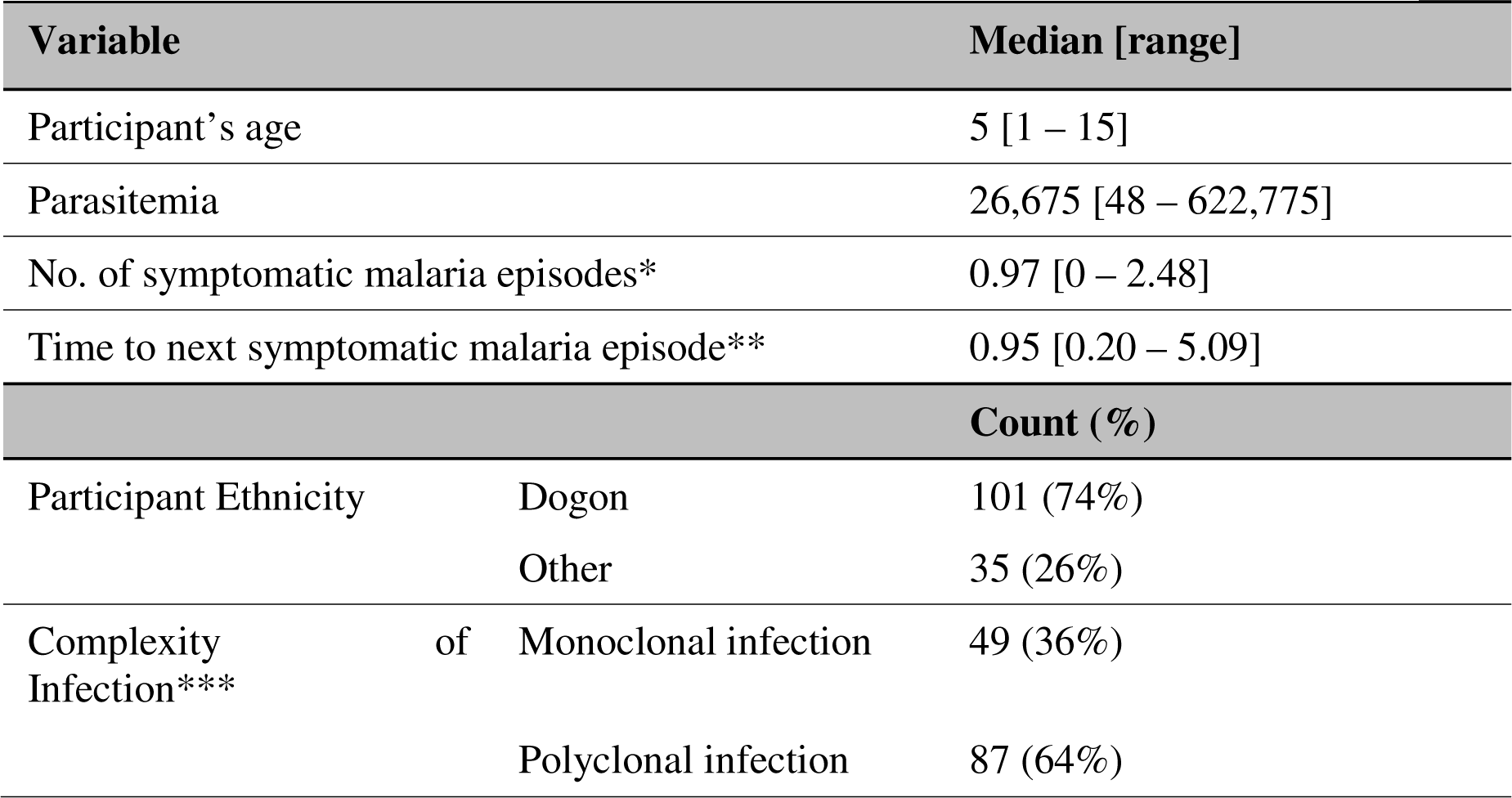
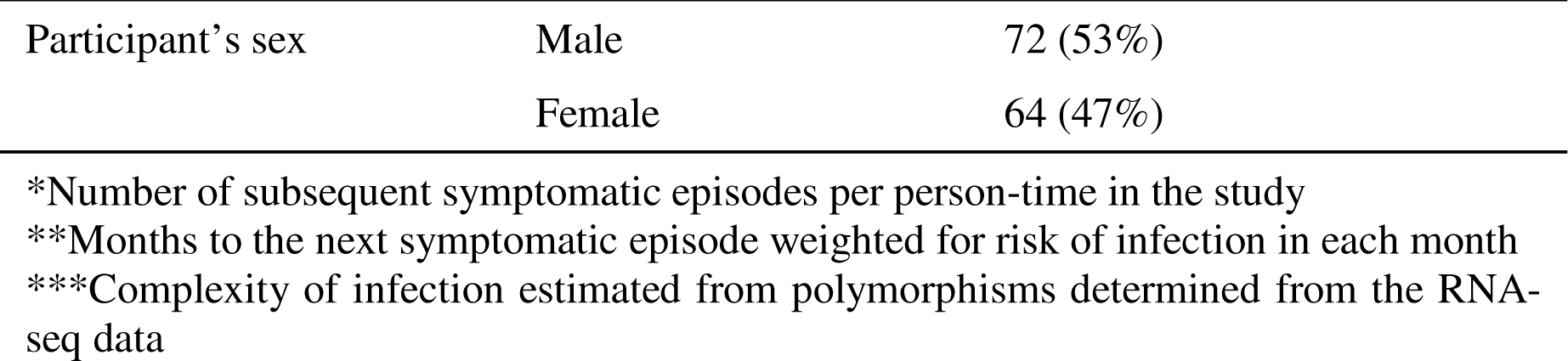
Characteristics of study participants.

| Variable |  |  | Median [range] |
| --- | --- | --- | --- |
| Participant's age |  |  | 5 [1 – 15] |
| Parasitemia |  |  | 26,675 [48 – 622,775] |
| No. of symptomatic malaria episodes* |  |  | 0.97 [0 – 2.48] |
| Time to next symptomatic malaria episode** |  |  | 0.95 [0.20 – 5.09] |
|  |  |  | Count (%) |
| Participant Ethnicity | Dogon |  | 101 (74%) |
|  | Other |  | 35 (26%) |
| Complexity of Infection*** | Monoclonal infection |  | 49 (36%) |
|  | Polyclonal infection |  | 87 (64%) |

|  |  |  |
| --- | --- | --- |
| Participant's sex | Male | 72 (53%) |
|  | Female | 64 (47%) |

\*Number of subsequent symptomatic episodes per person-time in the study
\*\*Months to the next symptomatic episode weighted for risk of infection in each month
\*\*\*Complexity of infection estimated from polymorphisms determined from the RNA-seq data

From each whole blood sample (N = 136), we extracted and sequenced RNA to characterize the host and parasite gene expression profiles (**Supplemental Table 1)**. To confirm that *P. falciparum* was responsible for each malaria episode and exclude possible co-infections, we first mapped all reads, simultaneously, to the genomes of *P. falciparum*, *P. malariae*, *P. ovale* and *P. vivax*. In each sample, 95% or more of the reads that mapped to a *Plasmodium* genome were uniquely mapped to *P. falciparum* and no co-infections were detected (**Supplemental Table 2**). (Note, that the small proportion of reads mapping to a species other than *P. falciparum* likely reflects reads derived from highly conserved regions that can be mapped to multiple species.)

We then mapped all reads simultaneously to the *P. falciparum* and human genomes (**Supplemental Table 1**) and obtained, on average, 85 million reads (21,121,310–149,135,348) mapped to the human genome (30.2% – 99.8%) and 14 million reads (160,712–64,001,204) mapped to the *P. falciparum* genome (0.2% – 69.8%) (**Supplemental Table 1**). The percent of reads mapped to the *P. falciparum* genome was significantly correlated with the parasitemia determined microscopically (p = 6.03 ×10^−14^, r^2^=0.34, **Supplemental Figure 1**). Overall, we were able to analyze variations in expression for 9,205 human and 2,484 *P. falciparum* genes.

We also leveraged the RNA-seq data to examine, within each infection, allelic variations at SNPs located in expressed parasite transcripts (*17, 19*) and determined that 87 of the 136 infections (64%) were polyclonal (**Supplemental Table 1**, see **Materials and Methods** for details).

### Variations in host and parasite gene expression are primarily driven by the child’s age and the infection’s parasitemia

To understand the contributions of different parameters to variations in host and parasite gene expression during symptomatic malaria episodes, we estimated the proportion of the variance in gene expression explained by the clinical and epidemiologic variables described in **Table 1**, considering all variables simultaneously (*20*). Overall, most of the variance in host and parasite gene expression was caused by inter-individual differences in gene expression (labeled “residuals” in our model).

From the factors we examined, only two variables contributed to the overall variance in host gene expression: the child’s age explained on average 3% of the overall variance in host gene expression (and between 0% and 39% of the variance of individual genes), while the infection’s parasitemia explained on average 2% (range = 0% to 23%) (**Figure 1A**). The remaining variables - the number of subsequent infections, the time to the next infection, the complexity of infections, or the sex of the participant – contributed very little to the overall variance in host gene expression (**Figure 1A**). Interestingly, although sex differences in the clearance of *P. falciparum* infections have been described (*21*) we did not observe significant sex differences in our data, with the exception of a small number of genes located on the sex chromosomes whose expression was significantly impacted by the sex of participants (**Supplemental Table 2**).

**Figure 1:**
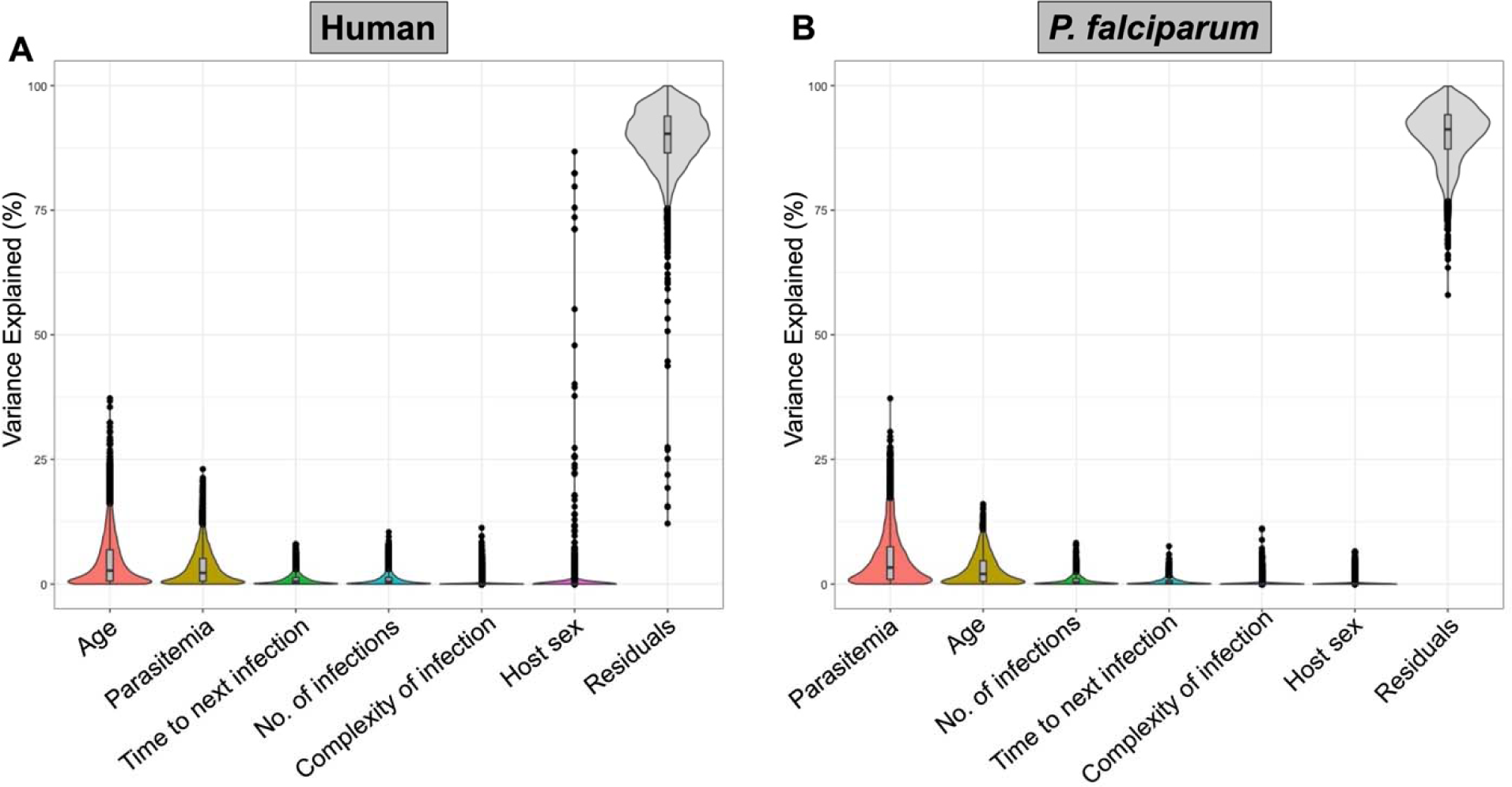
Percentage of the variance in host and parasite gene expression explained by each variable considered. Each violin plot shows the percentage of variance (y-axis) explained by each variable (x-axis) for each human (**A**) and *P. falciparum* (**B**) gene. Each black dot represents one gene, and the internal boxplot shows the mean and 25^th^ and 75^th^ percentiles. “Residuals” indicates the percentage of the variance in gene expression not explained by any of the variables considered (i.e., driven by remaining inter-individual differences).

Similarly, the variance in *P. falciparum* gene expression was partially explained by the parasitemia of the infection (median = 3%, range = 0% to 37%) and the child’s age (median = 2%, range = 0% to 16%), with the remaining variables explaining very little of the gene expression variance (**Figure 1B**).

We then statistically tested which specific host and parasite genes were differentially expressed according to these variables, accounting for the child’s sex and the month of the infection and correcting for multiple testing by false discovery rate.

Consistent with the results of the analysis of variance presented above, many host and parasite genes were significantly correlated with the child age and the infection’s parasitemia, while the number of subsequent infections and time to the next infection, the complexity of infection, or the sex of the participant were associated with only a small number of genes (**Table 2**, **Supplemental Table 3**, **Supplemental Table 5**).

**Table 2:** Differential gene expression. The table shows the number of human and *P. falciparum* genes associated with each clinical and epidemiologic variable (FDR = 0.1), before (unadjusted) and after (adjusted) adjusting for human immune cell or parasite developmental stage composition. All models were adjusted for the host sex and month of infection, and for age and parasitemia where appropriate.

| Variable |  | No. Human DEGs* | No. *P. falciparum* DEGs* |
| --- | --- | --- | --- |
| Parasitemia | Unadjusted | 3,221 | 1,779 |
|  | (Adjusted**) | (21) | (71) |
| Participant's age | Unadjusted | 4,174 | 833 |
|  | (Adjusted**) | (1,485) | (6) |
| No. symptomatic malaria episodes | - | 13 | 6 |
| Time to next symptomatic malaria episode | - | 0 | 0 |
| Complexity of infection | - | 0 | 0 |
| Participant's sex | - | 68 | 1 |

\*DEGs = differentially expressed genes (FDR = 0.1)
\*\*Adjusted for host immune cell or parasite developmental stage composition

### Host gene expression associated with parasitemia is mainly driven by differences in the proportions of neutrophils and T cells

3,221 human genes were associated with the infection’s parasitemia, after adjusting for the host’s age, sex, and the month of the infection (**Table 2**, **Figure 2A**). Many of the genes whose expression was positively correlated with parasitemia were neutrophil surface markers (e.g., CD177 (*22*)), granule/secretory vesicle proteins (e.g., MMP8 (*23*), MMP9 (*24*), ARG1 (*25*)), and genes involved in neutrophil recruitment (e.g., CXCR1 (*26*), CCRL2 (*27*)) (**Supplemental Table 3**). Conversely, the expression level of many genes related to T cells (e.g., CD3 (*28*), CD4, CD8 (*29*), and CXCR5 (*30*)) were negatively associated with parasitemia (**Supplemental Table 3**).

**Figure 2:**
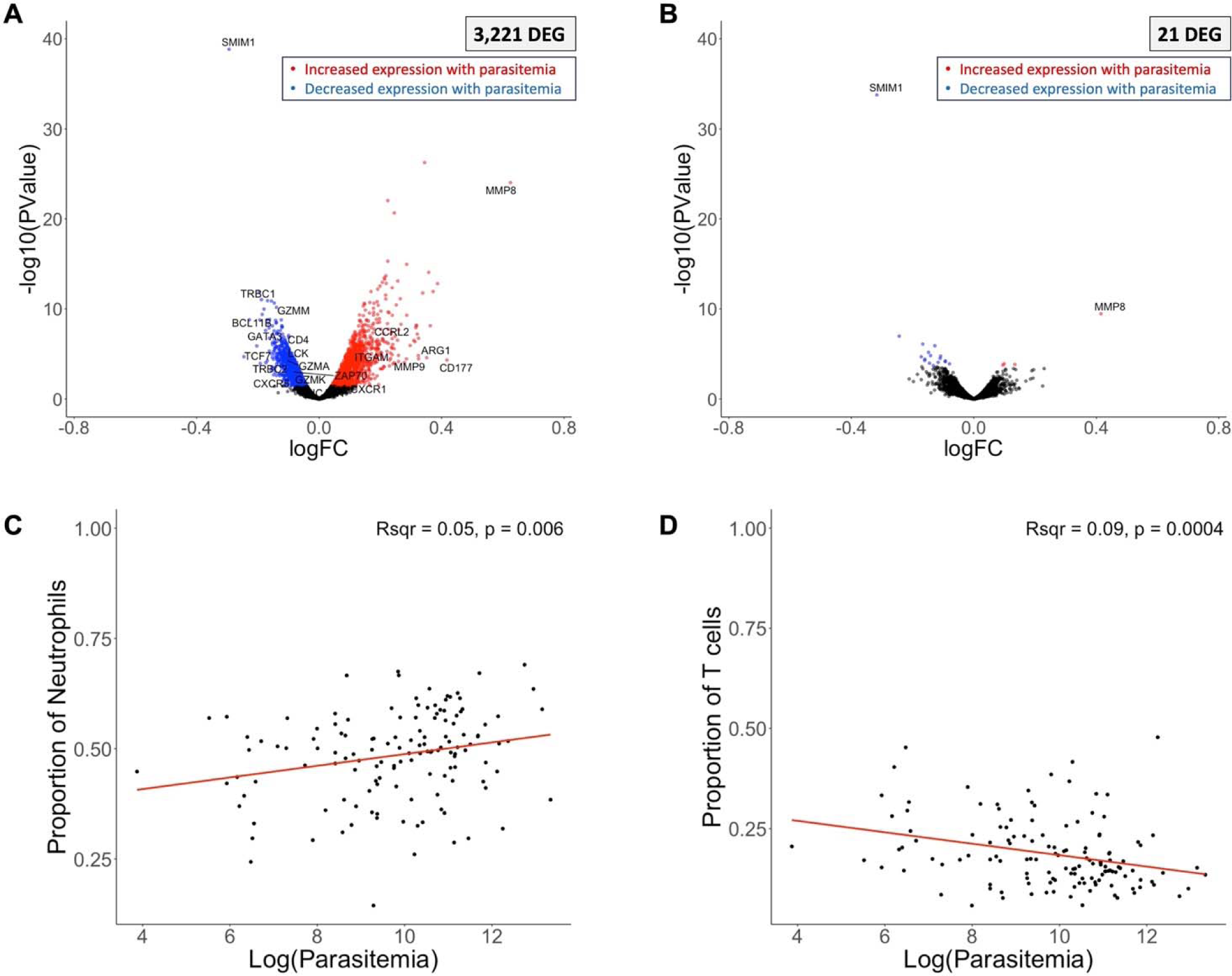
Host gene expression and parasitemia. The volcano plots show the association of host gene expression with the log of the parasitemia before **(A)** and after **(B)** adjusting for immune cell composition. Each point represents one gene, displayed according to its p-value (y-axis) and log fold-change (x-axis). Blue and red points represent genes that were significantly (FDR=0.1) more expressed in low and high parasitemia infections, respectively. Correlation of the proportion of neutrophils **(C)** or T cells **(D)** (y-axis), estimated by gene expression deconvolution, with the log of the parasitemia (x-axis). *DEG = differentially expressed gene.

To determine whether these differences in gene expression associated with parasitemia were i) genuine differences in gene regulation or ii) the result of differences in cell composition between samples, we adjusted the gene expression analyses for the relative proportion of each major immune cell subset determined, for each sample, from the RNA-seq data by gene expression deconvolution (*12*) (**Supplemental Table 4**). After correction for differences in cell composition, only 21 of the initial 3,221 differentially expressed host genes remained significantly associated with parasitemia (**Figure 2B**), suggesting that gene expression differences associated with parasitemia were mainly driven by heterogeneity in immune cell types, rather than differences in gene regulation. Two genes stood out in this analysis: SMIM1 and MMP8. SMIM1 is the surface marker for the Vel blood group (*31*) and is an understudied red blood cell (RBC) surface marker that may regulate RBC formation and hemoglobin concentration (*31*). SMIM1 has not been previously reported in malaria studies, but the correlation between its expression and parasitemia could indicate that it plays a role in regulating parasite development, possibly by modulating the amount of available hemoglobin (an important nutrient source for the parasites). Expression of MMP8, a neutrophil granule protein (*23*), has been shown to be elevated in the serum of individuals with uncomplicated malaria (*32*) and has been associated with malaria severity, particularly with cerebral malaria (*8, 33*). Although none of the individuals included in our study experienced cerebral malaria, it is possible that individuals with high parasitemia infections (and higher expression of MMP8) experienced more severe symptoms (symptom severity was not measured precisely in this cohort). Alternatively, this pattern of MMP8 expression could suggest a parasitemia-dependent neutrophil regulation, which warrants further study, particularly as it relates to immunopathology induced by this enzyme and its impact on disease severity.

We then examined which immune cell types were specifically associated with parasitemia. Consistent with previous reports (*34, 35*), we found that the relative proportion of neutrophils was significantly associated with parasitemia, with high parasitemia infections displaying a greater proportion of neutrophils (R^2^ = 0.05, p = 0.006) (**Figure 2C**). Neutrophils are important first responders in the innate immune system (*36, 37*) and have been reported to interact with *Plasmodium*-infected RBCs (*38, 39*) through phagocytosis and NET formation (*36–40*). Our results could indicate that neutrophils are released from the bone marrow into the peripheral blood proportionally to the number of parasites present, as circulating neutrophils attempt to combat the infection or, alternatively, that high parasitemia infections are characteristic of children with less developed immunity, relying more on a strong innate response (note, however, that not all children with high parasitemia infections were young (**Supplemental Figure 2**)).

Conversely, we found that the relative proportion of T cells was negatively associated with parasitemia: low parasitemia infections had, proportionally, more T cells than high parasitemia infections (R^2^ = 0.09, p = 0.004) (**Figure 2D**). Several non-exclusive mechanisms could explain this finding: i) T cell-mediated reduction of parasitemia (or initial reduction of hepatocyte infection, inhibiting blood stage development, which cannot be measured in this study), ii) parasite-mediated T cell inhibition, iii) lack of T cell stimulation at low parasitemia, and/or iv) T cell extravasation into secondary lymphoid tissues (so that they are missed in our peripheral blood samples).

Due to the relatively limited resolution of gene expression deconvolution that may hamper an accurate estimation of rarer cell types (*41, 42*), and to prevent data overfitting, we initially chose to conservatively estimate the proportion of only eight broadly-defined WBC subsets: neutrophils, T cells, B cells, mast cells, eosinophils, monocytes, NK cells and plasma cells. However, to assess whether a specific T cell subset was driving the correlation with parasitemia, we reiterated our gene expression deconvolution analysis and estimated the relative proportion of 22 immune cell subtypes included in our validated reference dataset, including seven different T cell populations (*13*) (**Supplemental Table 4**). We found that the proportion of naïve CD4 T cells (p = 0.00012, R^2^ = 0.10) and regulatory T cells (Treg) (p = 1.55 × 10^−5^, R^2^ = 0.13) were negatively correlated with parasitemia, while activated memory CD4 T cells (p = 0.013, R^2^ = 0.038) were positively correlated with parasitemia (**Supplemental Figure 3**). Some studies in mice have suggested that parasite-specific CD4 T cells directly reduce parasitemia (*43*). Our observed abundance of naïve CD4 T cells in low parasitemia infections in human children could indicate that, at the time of the blood collection, the parasitemia had already been controlled by the abundance of CD4 T cells. Tregs have been shown in human studies to have a complex and exposure-dependent role during *P. falciparum* infection and their relationship with parasitemia remains controversial (*44–46*). These cells expand after an immune response to modulate immunopathology induced by other cell types (*44*). These findings suggest either i) Treg expansion occurs after parasitemia has been controlled by other cell types (e.g., CD4 T cells) or ii) high parasitemia infections do not efficiently induce a Treg response, which could contribute to further immunopathology from these infections. Our observed enrichment for activated memory CD4 T cells in high parasitemia infections likely reflects the expansion of *P. falciparum*-specific memory cells in order to combat the infection in children who have developed some immunity from prior infections.

The analyses described above rely on estimations of the relative proportions of WBC subsets and the observations of more neutrophils and fewer T cells in high parasitemia infections might therefore not be independent. To better interpret these results, it will therefore be important to follow up on these observations with techniques such as flow cytometry that provides absolute quantitation and greater resolution. In addition, while blood samples were collected at the time that each individual presented to clinic with symptoms of malaria, it is possible that samples of different parasitemia were collected at different times after infection. This limitation (inherent to human field studies) could contribute to our observed differences in neutrophil and T cell proportions. Future work in animal models, where the infection and sampling times can be tightly controlled, could help to clarify the relationship between T cells and parasitemia, although important differences between animal models and human pathology exist (*47*).

Parasitemia and host age are not independent due to the gradual development of anti-malarial immunity with age and repeated exposures (*48–50*) (**Supplemental Figure 2**). Therefore, it is possible that, by adjusting the statistical analyses for age, we may have over-adjusted for genes that were correlated with both age and parasitemia. To attempt to address this confounding issue, we repeated our analyses, without correcting for age but using the largest possible subset of children in one narrow age range: four- and five-year-old children (N = 47). After correction for immune cell composition, we identified 143 genes associated with parasitemia (**Supplemental Figure 4B, Supplemental Table 6**), including many neutrophil effector proteins (e.g., MMP9 (*24*), LTF (*24*), PGLYRP1 (*51*)) and markers of neutrophil activation (e.g., CD177 (*22*), CD300H (*52*)) that displayed higher expression in high parasitemia infections. This result suggests that, in addition to the increase in circulating neutrophils, neutrophil regulation or proportions of different neutrophil subtypes (which were not sub-divided in our reference dataset) may also vary according to parasitemia, which should be investigated in more detail with higher resolution techniques such as flow cytometry. Similarly, we found that, in addition to a lower proportion of T cells, an important T cell growth factor (e.g., IL15 (*53*)) was negatively associated with parasitemia, suggesting differences in T cell activation at different levels of parasitemia (**Supplemental Table 6**). This finding further supports the hypothesis of an inefficient stimulation of T cells overall at low parasitemia and/or rapid control of parasitemia by memory cells before the time of sampling and warrants further investigation by basic immunology methods and flow cytometry in future cohorts.

Interestingly, we also detected multiple interferon (IFN) stimulated genes that were negatively associated with parasitemia, independently of age (**Supplemental Table 6**). While these genes can be expressed by a variety of cell types, this observation likely indicates that i) efficient IFN signaling leads to a dramatic reduction in parasitemia and ii) there is a critical threshold of parasites required to trigger efficient IFN signaling. These findings further highlight the complexity and heterogeneity of the antimalarial immune response among children and warrant studies, with larger sample sizes, to fully disentangle its relationship with parasitemia, independently from the patient’s age.

Note that since we measured gene expression correlated with parasitemia as a continuous variable, the log fold-change reflects the change in expression of each gene with each unit of parasitemia, which can be smaller than typical log fold-change values that measure differences in expression between two groups.

### Parasite gene expression associated with parasitemia is primarily driven by differences in stage composition

We identified 1,675 *P. falciparum* genes associated with parasitemia (**Figure 3A, Supplemental Table 5**). To determine whether these differences in gene expression were due to differences in developmental stage composition among samples, we estimated the relative proportion of each developmental stage in each sample by gene expression deconvolution (*15*) (**Supplemental Table 4**). After adjusting for stage composition, only 71 genes remained significantly associated with parasitemia (**Figure 3B**) including known antigens (e.g., PfHRP2 (*54*)), genes involved in DNA replication (e.g., PfRAD50 (*55*)), parasite asexual replication (e.g., PfACT1 (*56*)) and gametocyte development (e.g., PfNOT1-G (*57*), PfFACT-L (*58*)), which were positively associated with parasitemia, and genes involved in asexual development (e.g., PfCDPK1 (*59, 60*), PfPIC1 (*61*), PfHECT1 (*62*)), erythrocyte surface remodeling (e.g., PfFIKK1 (*63, 64*), PfFIKK7.1 (*63, 64*), PfSBP1 (*65*)) and gametocytogenesis (e.g., PfCDPK1 (*66*), PfPfs16 (*67*), PfMCA2 (*68*)), which were negatively associated with parasitemia (**Supplemental Table 5**).

**Figure 3:**
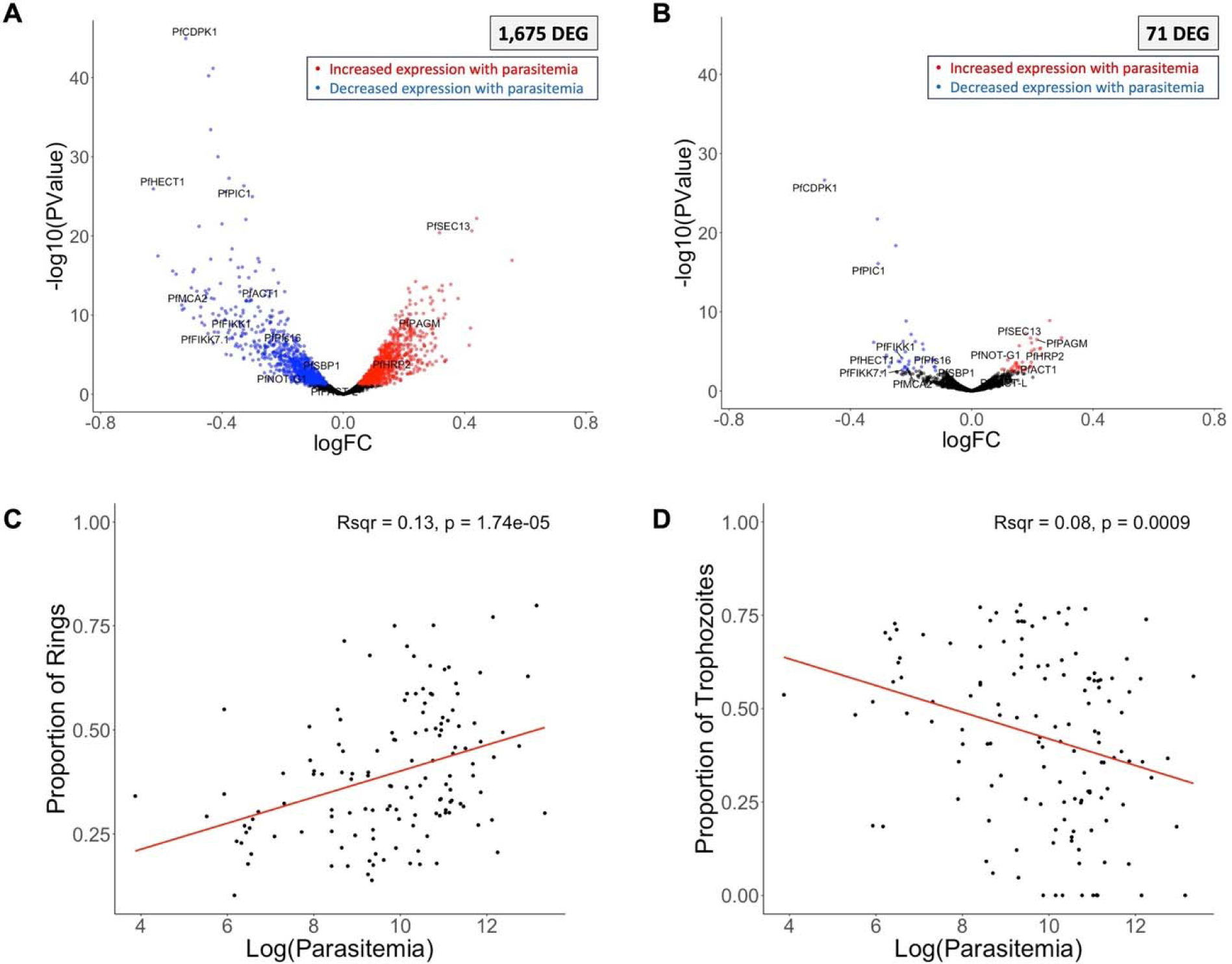
Parasite gene expression and parasitemia. The volcano plots show parasite gene expression associated with the parasitemia, unadjusted **(A)** and adjusted **(B)** for developmental stage composition. Each point represents one gene, displayed according to its p-value (y-axis) and log fold-change (x-axis). Blue and red points represent genes that were significantly (FDR=0.1) more expressed in low and high parasitemia infections, respectively. Correlation of the proportion of rings **(C)** or trophozoites **(D)** (y-axis), estimated from gene expression deconvolution, with the log of the parasitemia (x-axis). *DEG = differentially expressed gene.

We then tested which developmental stages varied with parasitemia. The relative proportion of ring stage parasites was positively correlated with parasitemia (R^2^ = 0.13, p = 1.74 × 10^−5^) (**Figure 3C**), while the relative proportion of trophozoite stage parasites was negatively correlated with parasitemia (R^2^ = 0.08, p = 0.0009) (**Figure 3D**).

Mature asexual *P. falciparum* parasites typically sequester in the tissues of infected patients and ring-stage parasites largely predominate in the peripheral blood (*69–71*). However, since ring-stage *Plasmodium* parasites are less transcriptionally active than other developmental stages (*72*), gene expression data can overestimate the relative proportion of mature stages (but proportionally in all samples (*15*)). The observed differences in developmental stage composition associated with parasitemia could therefore suggest that i) mature parasite sequestration is more efficient at high parasitemia and/or ii) that the regulation of intraerythrocytic development is parasitemia-dependent. Consistent with the latter hypothesis, several genes involved in asexual development remained negatively associated with parasitemia after adjusting for cell composition (e.g., PfCDPK1 (*59, 60*), PfPIC1 (*61*), PfHECT1 (*62*)) (**Figure 3B**), possibly indicating that, when parasitemia is high, parasites downregulate key genes to slow down their growth rate. Mouse models of *P. berghei* have recently shown that systemic host inflammation can slow maturation of the asexual parasites, suggesting that this parasitemia-dependent growth regulation is host mediated (*73*). Our results support this model for the first time in humans: higher parasitemia infections lead to more inflammation (see human gene expression results above) and this inflammatory environment could possibly explain the differences in asexual stage regulation observed in the *P. falciparum* gene expression.

As with the human gene expression analysis, since parasitemia and age are correlated, we may have missed parasite genes associated with parasitemia by over-correcting our model. We therefore repeated our gene expression analyses with the subset of 47 four- to five-year-old children, described above, and identified 421 genes associated with parasitemia after adjusting for developmental stage composition (**Supplemental Figure 5, Supplemental Table 6**). Consistent with our findings from the full cohort, we found several genes involved in invasion (e.g., PfGBP130 (*74*), PfPK2 (*75*), PfTrx-mero (*76*)) and replication (e.g., genes involved in cell cycle progression and chromosome organization) to be negatively associated with parasitemia.

Interestingly, several of these additional genes associated with parasitemia are consistent with parasitemia-dependent host-pathogen interactions. The expression of PfHMGB1, a gene that promotes host TNFα secretion in mouse models (*77*), was negatively correlated with parasitemia, suggesting that parasites modulate the host inflammatory response to increase their survival. In addition, PfEH1 and PfEH2 (*78*) were also less expressed in high parasitemia infections. These enzymes degrade erythrocyte-derived lipid signaling molecules (*78*) thereby reducing endothelial activation and decreasing the expression of ICAM1, an important ligand for parasite sequestration (*70*). At high parasitemia, lower expression of these enzymes may maintain a high level of ICAM1 expression in the endothelium, allowing for more efficient sequestration of parasites within tissues (consistent with our stage composition analyses) and facilitating evasion of host immunity. These possible mechanisms of parasitemia-dependent modulation of host inflammation and parasite sequestration will need to be validated but provide exciting hypotheses for studying unexplored density-dependent host/pathogen interactions in malaria infections.

### Host gene expression associated with participant age is partially explained by differences in immune cell composition

We identified 4,174 genes with expression significantly associated with host age (which ranged from 1 to 15 years old in our cohort) (**Table 2**, **Figure 4A, Supplemental Table 3**). To determine whether these differences in gene expression were explained by cell type heterogeneity among samples, we examined the correlation between host age (as a continuous variable) and the relative proportion of each immune cell type. The proportion of neutrophils was positively associated (R^2^ = 0.07, p = 0.0019), and the proportions of B cells (R^2^ = 0.13, p = 2.10 × 10^−5^), NK cells (R^2^ = 0.13, p = 1.15 × 10^−5^) and plasma cells (R^2^ = 0.06, p = 0.004) were negatively associated with age (**Figure 4C-F**). In contrast to the gene expression differences associated with parasitemia, host gene expression associated with age was only partially explained by changes in cell composition: over one third of the differentially expressed genes (N = 1,485) remained significantly associated with age after adjusting for WBC composition (**Table 2**, **Figure 4B**).

**Figure 4:**
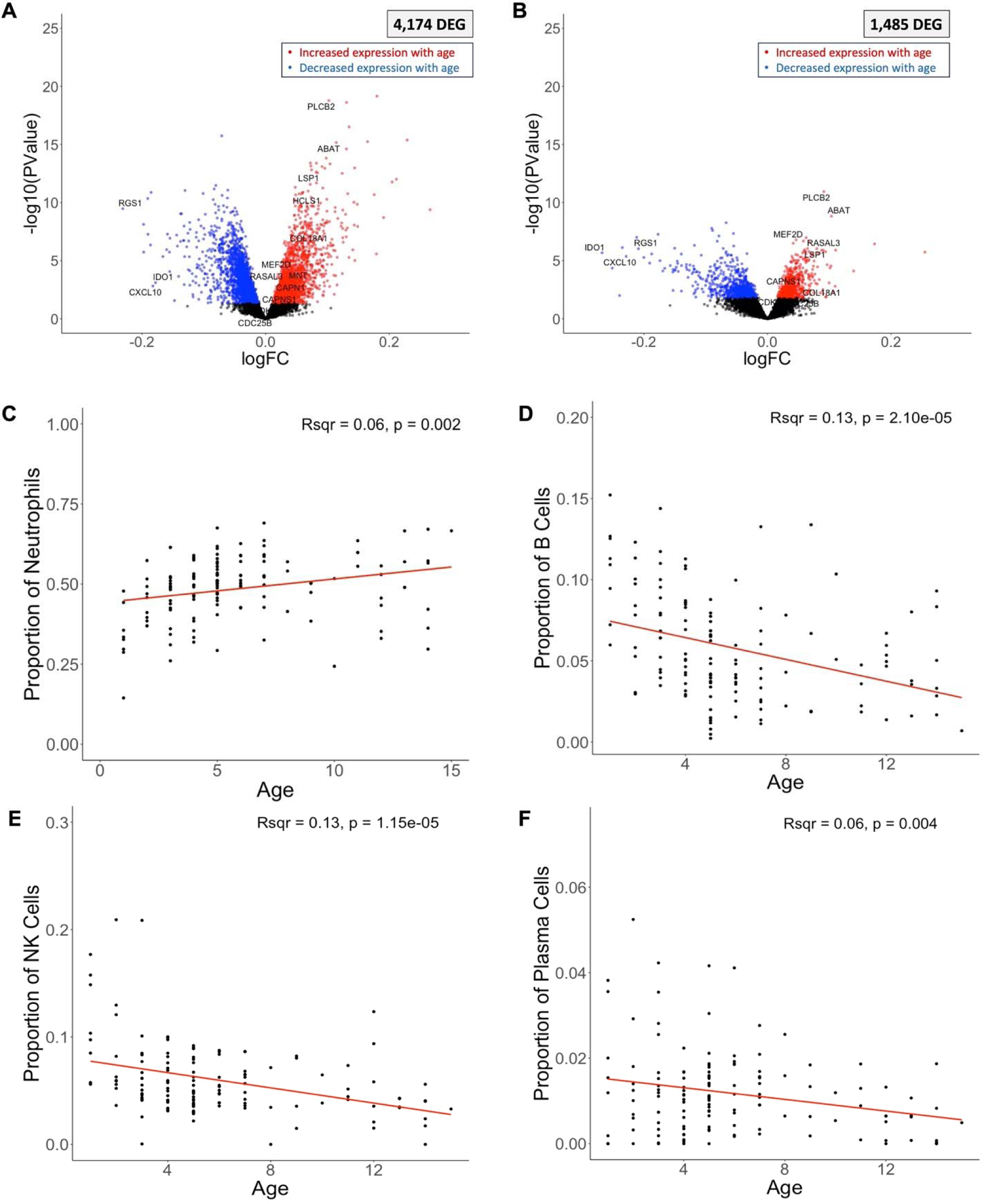
Host gene expression and child’s age. The volcano plots show the association between each human gene’s expression and the child’s age at the time of the infection, before **(A)** and after **(B)** adjusting for differences in immune cell composition. Each dot represents one gene and is displayed according to its log10 p-value (y-axis) and fold-change (x-axis). Blue and red points represent differentially expressed genes (FDR=0.1) that were more expressed in younger and older children, respectively. Correlation of the proportion of neutrophils **(C),** B cells **(D),** NK cells **(E),** and Plasma cells **(F)** (y-axis), estimated from gene expression deconvolution, with the age of the child in years (x-axis). Note that different ranges for the y-axis in C, D, E and F due to differences in cell proportions. *DEG = differentially expressed gene.

To contextualize the genes that remained associated with age after adjusting for cell composition, we used the KEGG database (*79–81*) to examine their distribution among key relevant pathways using Pathview (*82, 83*). The gene expression patterns in older children were consistent with activation of adaptive immunity, including activation of platelets (i.e., PLCB2, Pi3K) (**Supplemental Figure 6A**), T cell metabolism (i.e., ABAT (*84*), MEF2D (*85, 86*)), and neutrophil inflammatory response (i.e., RASAL3 (*87*), LSP1 (*88–90*)) (note that neutrophil activation is not represented in the KEGG database) (**Supplemental Table 3**). We further found increased expression of several genes involved in TCR (**Supplemental Figure 6B**) and BCR (**Supplemental Figure 6C**) signaling pathways.

While acquisition of immune memory to *P. falciparum* is complex and not entirely understood (*91–93*), taken together, this pattern is consistent with a T cell memory response. Neutrophil and platelet activation can enhance the adaptive immune response in general (*94–96*), as well as memory CD4 T cell responses, specifically (*94, 97, 98*). Memory CD4 T cell responses have also been observed upon repeated infections in mouse models (*99–101*). After re-infection, *Plasmodium*-specific memory CD4 T cells rapidly proliferate to respond to the pathogen. Indeed, we also observed increased expression of several genes involved in cell proliferation (*102*) (e.g., CAPN1, CAPNS1, CDC25B, CDK9, COL18A1, HCLS1, MNT) positively correlated with age (**Supplemental Table 3**).

Interestingly, several genes involved in the regulation of the actin cytoskeleton (**Supplemental Figure 6D**) and focal adhesion (**Supplemental Figure 6E**) pathways were also positively correlated with age, which could indicate immune synapse formation and/or leukocyte extravasation. Though not specific to memory lymphocytes, the immune synapse is required for activation of T and B cells (*103*). The actin cytoskeleton has also been shown to undergo remodeling after successful TCR (*104*) and BCR (*105*) signaling. Because activation of memory lymphocytes is faster than naïve lymphocytes (*106*), we speculate that the association of an activated adaptive response in older children at symptom presentation is consistent with a memory response after years of exposure and immune system aging.

The gene expression pattern in younger children was broadly suggestive of an innate inflammatory response, including genes whose expression is induced by interferon signaling (e.g., RGS1 (*107*), IDO1 (*108*), CXCL10 (*109*)), NOD-like receptor (NLR) signaling (**Supplemental Figure 7A**), Toll-like receptor (TLR) signaling (**Supplemental Figure 7B**), phagocytosis (**Supplemental Figure 7C**) and antigen presentation (**Supplemental Figure 7D**).

This innate-dominated immune environment in younger children (here, the lower limit of age being 1 year old with most children in the cohort being between 1 and 5 years old) is consistent with the clinical observation that immunological memory to malaria does not develop until later in adolescence and adulthood (*93*). These findings also likely reflect the maturation of the immune system over time, which has been shown to impact anti-malarial immunity (*49, 50*), but has been largely under studied in healthy children in this age range, particularly in Sub-Saharan African populations.

Both TLR and NLR signaling are key components of the innate response to *Plasmodium* infection (*110*). While it is still unclear how NLR signaling pathways impact anti-malarial immunity, hemozoin, a toxic byproduct of *Plasmodium* digestion of hemoglobin, can stimulate NLRP3 (an NLR) *in vitro* (*111*). TLR recognition of *Plasmodium* has been better characterized (*112*): TLR9 recognizes DNA-hemozoin complexes, TLR1/2 heterodimers recognize GPI anchors of *Plasmodium* proteins, and TLR7 and TLR8 recognize *Plasmodium* RNA. TLR and NLR signaling share a common result: the production of interferons, mediating host defense against the pathogen, as well as immunopathology (*112*).

Because phagocytosis is a key component of antigen presentation (*113*), these pathways are likely linked. Antigen presentation is a key step in bridging the innate and adaptive immune systems (*113*) and our findings suggest that the innate immune system in younger children is actively responding to *P. falciparum* at the time of sample collection (i.e., at symptom presentation).

Overall, our findings highlight that different immune pathways are preferentially activated upon *P. falciparum* infection depending on the age of the child, progressing from a greater reliance on innate immunity to acquired immunity as a child ages. These findings provide a potential mechanism underlying the gradual acquisition of immunity, first against severe disease and eventually from all malaria symptoms as children age, as has been reported in several epidemiological (*114, 115*) and clinical studies (*116*). While age in this context is likely to be at least partially a proxy for exposure to *P. falciparum,* as children in our cohort experience, on average, two malaria infections per transmission season (*18*), both age and repeated exposure have been independently linked to clinical protection from malaria and development of anti-*Plasmodium* immunity in Ugandan children (*48, 117*). Additionally, studies of Indonesian adults have shown an age-dependent reduction in risk of malaria disease, independent of prior exposures, suggesting that development of the immune system with age influences clinical protection (*49, 50*). Here, our results provide evidence for age-dependent gene expression in Malian children, which could be linked to the development of clinical protection from malaria and will be important to validate with basic immunology methods and larger cohorts in future studies. Additionally, future studies with access to prior infection history to precisely account for malaria exposure in each child, independent of age, will be important in disentangling gene expression separately associated with age and exposure history.

Again, because age and parasitemia are correlated, we may miss genes that were truly associated with age by correcting for parasitemia. We attempted to analyze subsets of children with similar parasitemia but of different ages by creating bins of parasitemia with a one log range, but the low sample sizes within each bin precluded meaningful analyses and further studies are necessary.

### The proportion of male gametocytes is associated with participant age

Out of 2,484 *Plasmodium* genes, 833 were associated with participant age (**Table 2**, **Figure 5A**, **Supplemental Table 5**). To determine whether these differences in gene expression were explained by differences in stage composition between samples, we adjusted for the relative proportion of each developmental stage (**Supplemental Table 4**). After adjusting, only six *P. falciparum* genes remained associated with participant age (**Table 2**, **Figure 5B**, **Supplemental Table 5**).

**Figure 5:**
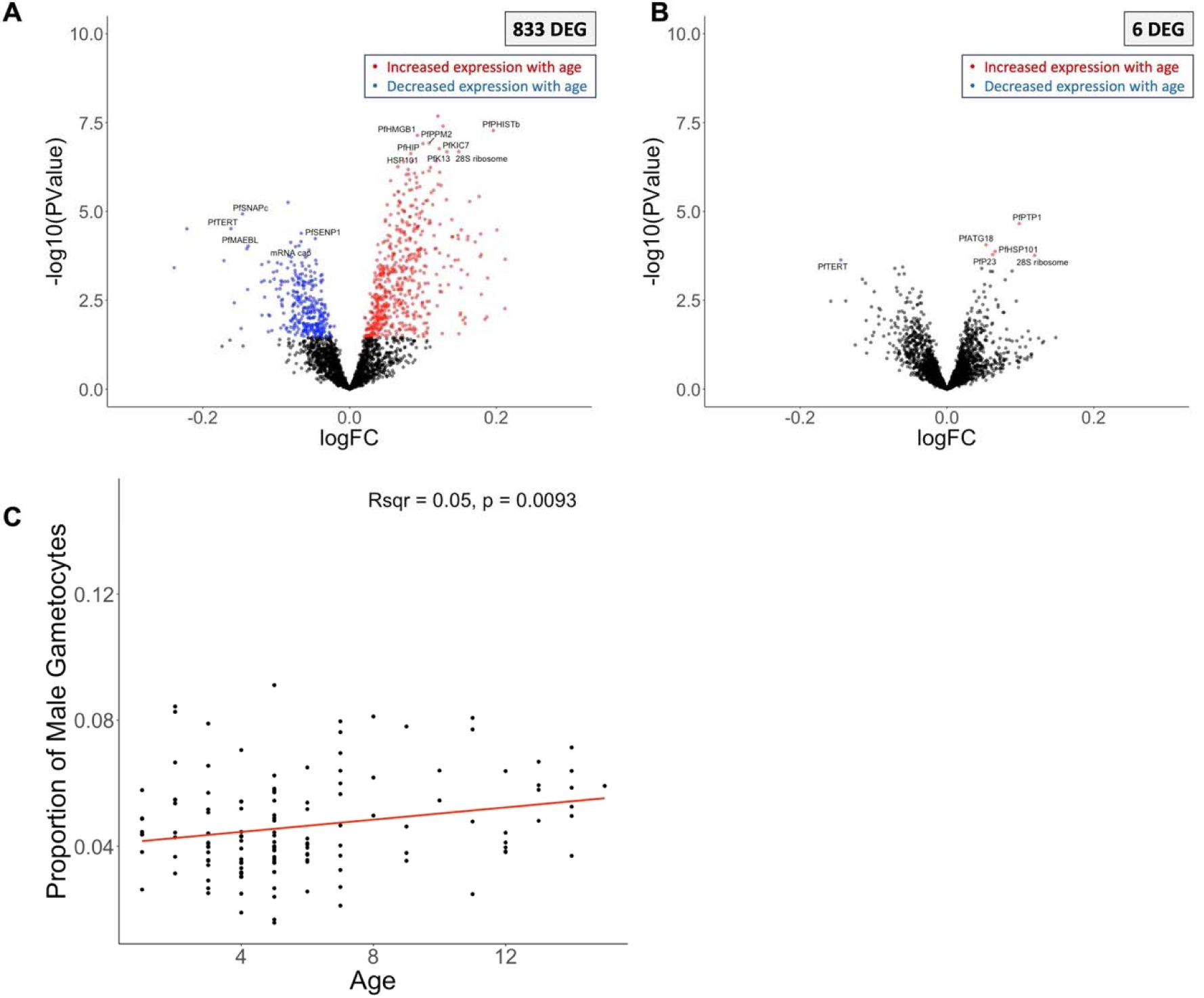
Parasite gene expression and child’s age. The volcano plots show parasite gene expression associated with the child’s age at the time of infection, before **(A)** and after **(B)** correction for developmental stage composition. Each point represents one gene, displayed according to its p-value (y-axis) and fold-change (x-axis). Blue and red points represent differentially expressed genes (FDR=0.1) that were more expressed in younger and older children, respectively. Correlation of the proportion of male gametocytes (y-axis) inferred from gene expression deconvolution with the participant’s age in years (x-axis) (**C**). Note that different ranges for the y-axis in C due to differences in cell proportions. *DEG = differentially expressed gene.

Interestingly, the proportion of male gametocytes was positively associated with participant age (R^2^ = 0.049, p = 0.0093) (**Figure 5C**). While gametocytogenesis and gametocyte development are not yet completely understood, *Plasmodium* parasites have been shown to vary their sex ratio according to environmental changes (*118, 119*), including host immune status (*118, 120*). Consistent with our findings, a few studies have also reported a male-skewed gametocyte sex ratio in older *P. falciparum*-infected children (*121, 122*). Since variation in gametocyte sex ratio is thought to impact transmission success (*119, 123*) and male-dominated ratios have been shown to increase transmission from infected humans to mosquitos (*123*), our observations could indicate that older children are more likely to contribute to disease transmission, consistent with previous epidemiology work from other settings with continuous, rather than seasonal transmission (*124, 125*). This result further highlights that malaria elimination initiatives should consider age (in addition to immunity status) when prioritizing interventions.

It is important to note that because malaria cases were detected by active case detection, it is possible that older children became symptomatic later after infection initiation than younger children due to increasing immunity with age (*48, 126*). “Older” infections are more likely to harbor gametocytemia, as *P. falciparum* gametocytes appear in the blood after about two weeks (*118*). However, while age of infection may contribute to gametocytemia overall, it is unlikely that this entirely explains the male-skewed gametocytemia we observed here.

### Genes related to a type 1 IFN response are associated with protection from future malaria episodes

While the number of subsequent symptomatic malaria episodes in the study period was not a major driver of the host gene expression, 13 genes were significantly associated this variable, including three genes that are characteristic of (although not exclusive to) a type 1 IFN (IFN1) response: CXCL10 (*109*), SOCS1 (*127*), PLAAT4 (*128*) (**Supplemental Figure 8, Supplemental Table 3**). The effects of IFN1 in response to malaria are variable and depend on both host genetic background and parasite genotype (*112*), but they have been shown to influence T cell activation (*129*) and antibody production (*130*). CXCL10 expression can also lead to growth acceleration of *P. falciparum in vitro* (*131*). Likewise, we found a positive correlation between CXCL10 with parasitemia, potentially suggesting a positive feedback-like interaction between the host and parasite, whereby parasites stimulate an IFN1 response in the host, leading to CXCL10 production, which can both increase parasitemia and modulate protection against future infections by influencing the adaptive immune response. Future mechanistic immunology studies are necessary to precisely disentangle this relationship.

### Gametocyte markers are associated with susceptibility to future malaria episodes

Similar to the host gene expression, few parasite genes were significantly associated with the number of subsequent symptomatic infections in the study period (N = 6) but those included those known regulators of gametocytogenesis (PfG27/25 and PfAP2-G) (**Supplemental Figure 9**) (**Supplemental Table 4**). This observation is interesting given previous reports linking higher gametocyte density with an anti-inflammatory environment (*132, 133*). This could suggest that failure to develop a sufficient inflammatory response to one infection could promote gametocyte production and reduce the development of long-term, protective immunity. Again, because gametocytes are the transmissible stage of *P. falciparum,* it is important to understand the interplay between susceptibility and gametocytogenesis to both protect susceptible children and prevent transmission of the parasites.

### Conclusions

Here, we described associations of host and parasite gene expression with several clinical and epidemiologic variables during symptomatic *P. falciparum* infections. Parasitemia and host age explained most of the variance in gene expression, while very few genes were associated with the complexity of infection, host sex, and the number of and time to the next malaria episode. Gene expression differences associated with parasitemia were almost entirely caused by differences in the proportions of host immune cells and parasite developmental stages among samples, while many genes remained significantly associated with age after accounting for cell composition. Our findings highlight that the host and parasite gene expression profiles are strongly influenced by differences in the inflammatory environment that are mediated by the parasitemia of the infection and the age of the host. Overall, this study provides new insights into the regulation of the human immune system and *P. falciparum* intraerythrocytic development during uncomplicated symptomatic infections and highlight the importance of rigorously considering the child’s age for targeted treatment and elimination strategies.

## MATERIALS AND METHODS

### Ethics approval and consent

Individual informed consent/assent was collected from all children and their parents. The study protocol and consent/assent processes were approved by the institutional review boards of the Faculty of Medicine, Pharmacy and Dentistry of the University of Maryland, Baltimore and of the University of Sciences, Techniques and Technologies of Bamako, Mali (IRB numbers HCR-HP-00041382 and HP-00085882).

### Samples

We selected 136 whole blood samples, collected directly in PAXgene blood RNA tubes, from children experiencing a symptomatic uncomplicated malaria episode caused by *Plasmodium falciparum* parasites. The presence of parasites and the parasite species were initially determined by light microscopy using thick blood smears. All infections were successfully treated with antimalarial drugs according to the Mali National Malaria Control Programme standards.

### Case definition

Children were classified, by the field clinicians, as experiencing symptomatic uncomplicated malaria if they i) sought treatment from the study clinic, ii) experienced symptoms consistent with malaria (i.e., fever, headache, joint pain, abdominal pain, vomiting or diarrhea), and iii) *Plasmodium falciparum* parasites were detected, at any density, by thick blood smear, and if they lacked any signs of severe malaria (e.g., coma, seizures, severe anemia) (*18*).

### Weighted number of and time between subsequent infections

To correct for the variation in risk of transmission throughout the study period, we weighted the number of, and time between, infections for the relative risk of malaria during each child’s person-time in the study. To calculate the number of subsequent infections, we first calculated, for the whole cohort, the number of malaria cases per month divided by the total number of children followed-up in that month (month-weight). Next, we calculated each child’s person-time left in the study after our sequenced infection by taking the sum of the month-weights for each month during which a child was enrolled after the date of our sequenced infection. Finally, we divided each child’s number of symptomatic malaria episodes occurring after our sequenced infection by their person-time remaining. To calculate the time to the next subsequent symptomatic malaria episode for each child, we summed the month-weights for each month between our sequenced infection and the next documented symptomatic malaria episode.

### Generation of RNA-seq data

We extracted RNA from whole blood using MagMax blood RNA kits (Themo Fisher). Total RNA was subjected to rRNA depletion and polyA selection (NEB) before preparation of stranded libraries using the NEBNext Ultra II Directional RNA Library Prep Kit (NEB). cDNA libraries were sequenced on an Illumina NovaSeq 6000 to generate ∼55-130 million paired-end reads of 75 bp per sample. To confirm that *P. falciparum* was responsible for each malaria episode, we first aligned all reads from each sample using hisat2 v2.1.0 (*134*) to a fasta file containing the genomes of all *Plasmodium* species endemic in Mali downloaded from PlasmoDB (*135*) v55: *P. falciparum* 3D7*, P. vivax* PvP01*, P. malariae* UG01, and *P. ovale curtisi* GH01. After ruling out coinfections and misidentification of parasites, we aligned all reads using hisat2 to a fasta file containing the *P. falciparum* 3D7 and human hg38 genomes i) using default parameters and ii) using (--max-intronlen 5000). Reads mapping uniquely to the hg38 genome were selected from the BAM files generated with the default parameters. Reads mapping uniquely to the *P. falciparum* genome were selected from the BAM files generated with a maximum intron length of 5,000 bp. PCR duplicates were removed from all files using custom scripts. We then calculated read counts per gene using gene annotations downloaded from PlasmoDB (*P. falciparum* genes) and NCBI (human genes) and the subread featureCounts v1.6.4(*136*).

### Gene expression analysis

Read counts per gene were normalized into counts per million (CPM), separately for human and *P. falciparum* genes. To filter out lowly expressed genes, only human or *P. falciparum* genes that were expressed at least at 10 CPM in > 50% of the samples were retained for further analyses (9,205 and 2,484 genes, respectively). Read counts were normalized via TMM for differential expression analyses. Statistical assessment of differential expression was conducted, separately for the human and *P. falciparum* genes, in edgeR (v 3.32.1) (*137*) using a quasi-likelihood negative binomial generalized model i) with and without correcting for proportion of the major human immune cell types for human genes and ii) with and without correcting for proportion of each parasite developmental stage for *Plasmodium* genes. Age and parasitemia were considered as continuous variables using edgeR (*137*). We log-transformed parasitemia estimates to fit the data to a normal distribution. Models used to estimate the gene expression associated with parasitemia were adjusted for host age, month of infection and host sex. Models used to estimate the gene expression associated with host age were adjusted for parasitemia, month of infection and host sex. All results were corrected for multiple testing using FDR (*138*).

### Gene expression deconvolution

CIBERSORTx (*12*) was used to estimate, in each sample, the proportion of i) human immune cell types and ii) *Plasmodium* developmental stages, separately, directly from the RNA-seq data. To deconvolute human gene expression profiles, we used as a reference LM22 (*13*), a validated leukocyte gene signature matrix which uses 547 genes to differentiate 22 immune subtypes (collapsed to eight categories in our analysis to prevent data sparsity). A custom signature matrix derived from *P. berghei* scRNA-seq data (*139*) was used for *P. falciparum* stage deconvolution, using orthologous genes between the two species (*15*).

### Complexity of infection

We used GATK GenotypeGCVFs (*140*) to call variants in all samples directly from the RNA-seq reads and analyze the complexity of each infection examined. While this pipeline was initially developed for analyzing whole genome sequence data, we previously showed that it can be applied to RNA-seq data (*17*). Briefly, we filtered the genotype file to retain only positions that had a maximum of two alleles, no more than 20% missing information and to remove positions within *Plasmodium* multi-gene families (due to inaccurate mapping of reads within these regions because of high sequence variability) (*17*). To determine the complexity of each infection (i.e., monoclonal vs. polyclonal), we then estimated F_ws_ from the filtered genotype file from GATK using moimix (*141*). Samples with F_ws_ > 0.95 were considered monoclonal and F_ws_ < 0.95 polyclonal.

## Acknowledgements

We thank the participants and their families for participating in this study, as well as the community of Bandiagara, Mali.

## Funding

National Institutes of Health grant R21AI146853 (DS, MAT)

National Institutes of Health grant R01HL146377 (MAT)

National Institutes of Allergy and Infectious Diseases pre-doctoral fellowship T32 AI095190 (KT)

National Institutes of Health grant U01AI065683 (CVP)

National Institutes of Health grant R01HL130750 (CVP)

National Institutes of Health grant R01AI099628 (MATh)

National Institutes of Health grant D43TW001589 (CVP)

## Author contributions

Conceptualization: KT, MAT, DS

Methodology: KT, DS, MAT

Investigation: KT, DS

Visualization: KT

Funding acquisition: KT, DS, MAT, CVP, MATh

Project administration: SY, DC, AK, AD, YT, KT, AN, BK, OD

Supervision: DS

Writing – original draft: KT, DS

Writing – review & editing: KT, DS, MAT, ML, ES, AB, KEL, STH

## Competing interests

Authors declare that they have no competing interests.

## Data and materials availability

All sequence data generated in this study are deposited in the Sequence Read Archive under the BioProject PRJNA962942. Custom scripts are available at https://github.com/tebbenk/symptomatic_malaria.

## Notes

### Competing Interest Statement

The authors have declared no competing interest.

