## Supplemental Table Legends for "Gene expression analyses reveal differences in children’s response to malaria according to their age"

*All tables uploaded as excel files

**Supplemental Table 1:** Sequencing statistics and clinical variables for each individual. Number of reads mapped to each *Plasmodium* species for each individual.

**Supplemental Table 2:** Proportion of the variance in expression of each human and *Plasmodium* gene explained by each of the variables in Table 2.

**Supplemental Table 3:** Human gene expression correlated with each variable presented in Table 2, both unadjusted and adjusted for cell composition.

**Supplemental Table 4:** Proportion of human immune cells and *P. falciparum* developmental stages in each individual estimated by CIBERSORTx.

**Supplemental Table 5:** *P. falciparum* gene expression correlated with each variable presented in Table 2, both unadjusted and adjusted for cell composition.

**Supplemental Table 6:** Human and *P. falciparum* gene expression correlated with parasitemia at a subset of children aged four to five years old, adjusted for cell composition.

**Supplemental Table 7:** Human gene expression associated with age in subsets of children with similar parasitemia, unadjusted and adjusted for immune cell composition.
