## Supplemental Figures for "Gene expression analyses reveal differences in children’s response to malaria according to their age"

**Supplemental Figure Legends:**

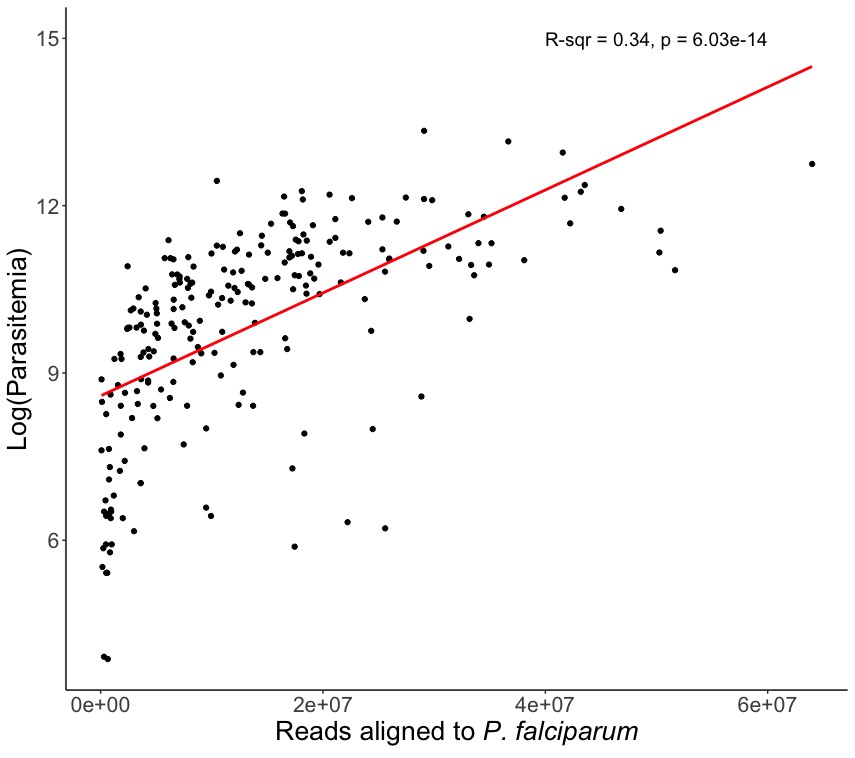

**Supplemental Figure 1: Parasitemia and mapped reads are highly correlated.** Log of *P. falciparum* parasitemia measured by microscopy (y) vs. number of reads mapping to *P. falciparum* (x).

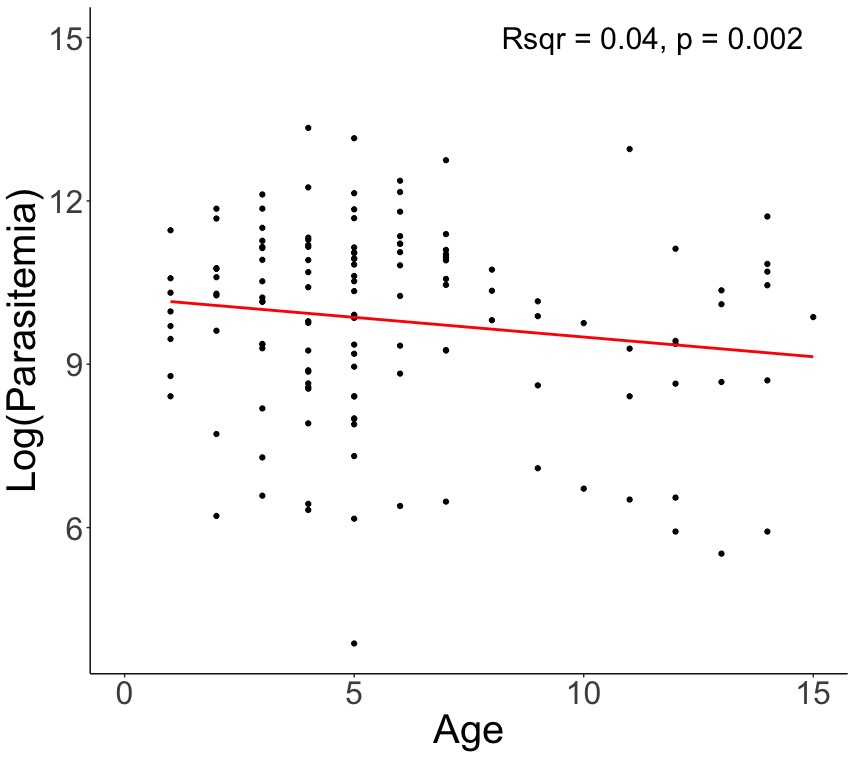

**Supplemental Figure 2: Age and parasitemia are weakly correlated, with older children having lower parasitemia.** Age in years (y) vs. the log of the *P. falciparum* parasitemia measured by microscopy.

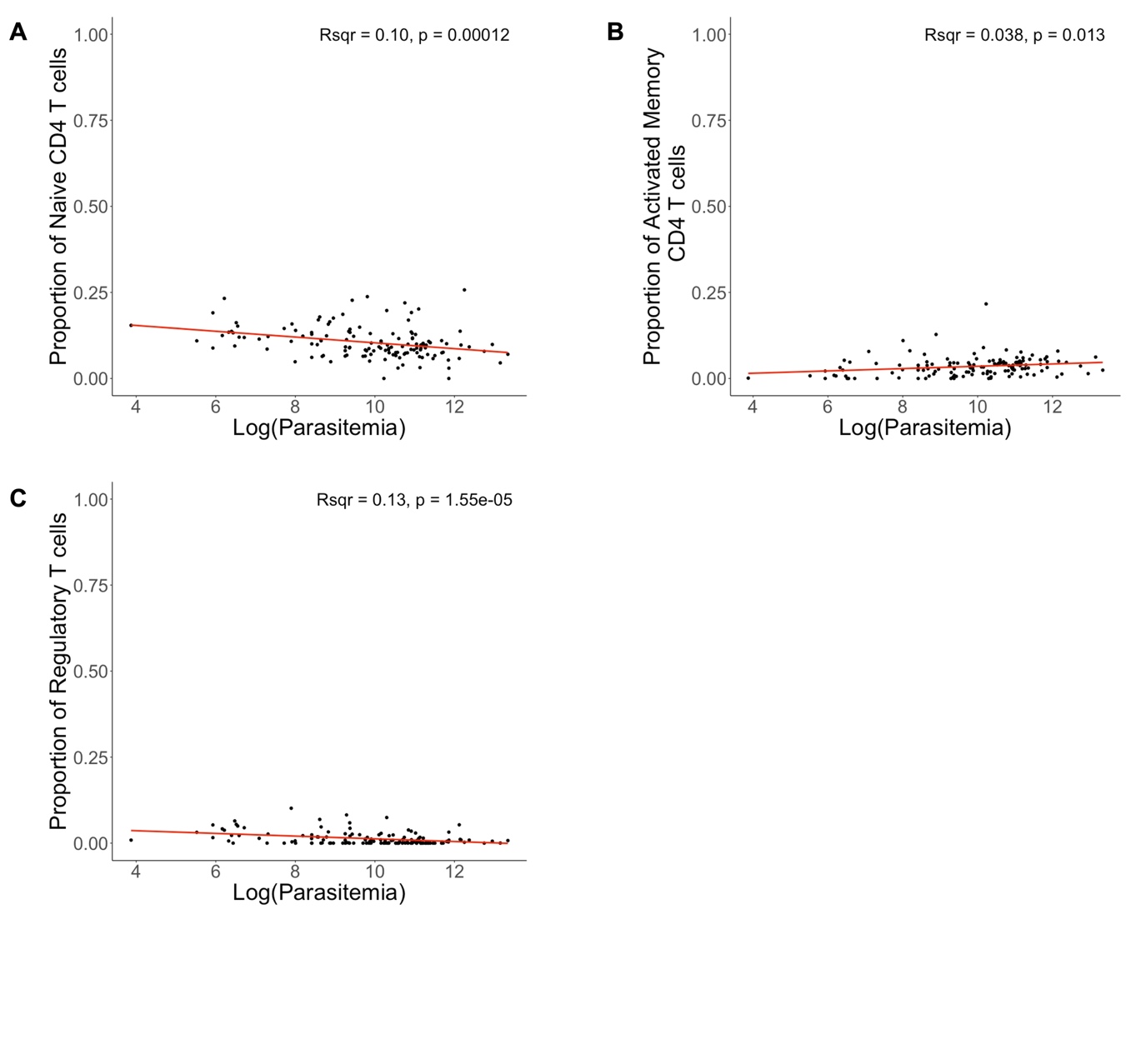

**Supplemental Figure 3: The proportion of specific T cell subsets, estimated by gene expression deconvolution that are correlated with the log of the *P. falciparum* parasitemia measured by microscopy. A)** Proportion of naïve CD4+ T cells correlated with log(parasitemia) **B)** Proportion of activated CD4+ memory T cells correlated with log(parasitemia) **C)** Proportion of regulatory T cells correlated with log(parasitemia).

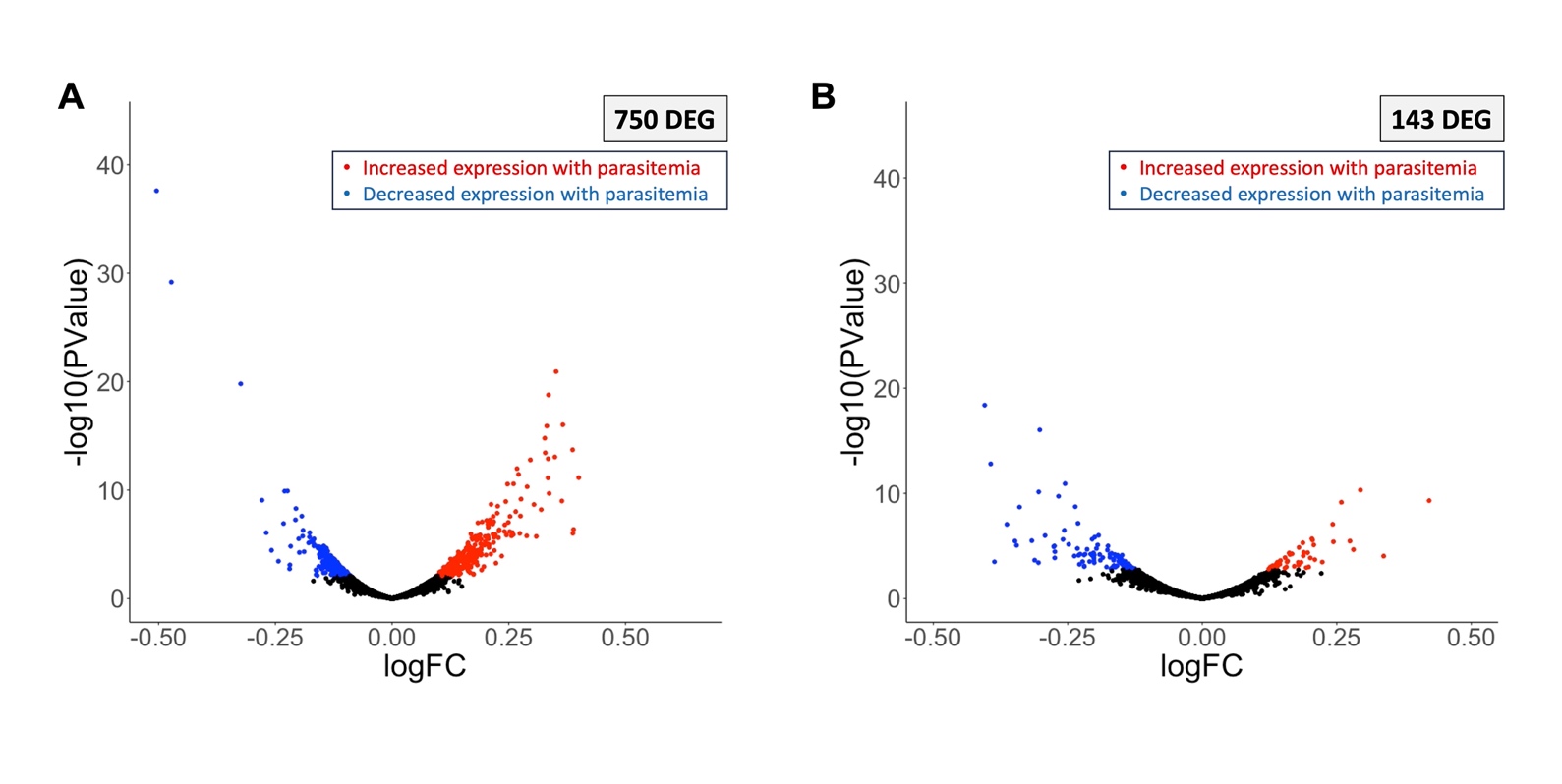

**Supplemental Figure 4: Some human genes are correlated with parasitemia, separate from age.** Volcano plot of human genes associated with log(parasitemia) in four- to five-year-old children. Each point represents one gene, plotted by its P-value (y) and log(fold-change) value (x). Blue points are genes that are significantly negatively associated with log(parasitemia). Red points are genes that are significantly positively associated with log(parasitemia). **A)** Model unadjusted for immune cell composition; 750 significant genes. **B)** Model adjusted for immune cell composition; 143 significant genes. Significantly differentially expressed genes have an (FDR = 0.1). *DEG = differentially expressed gene.

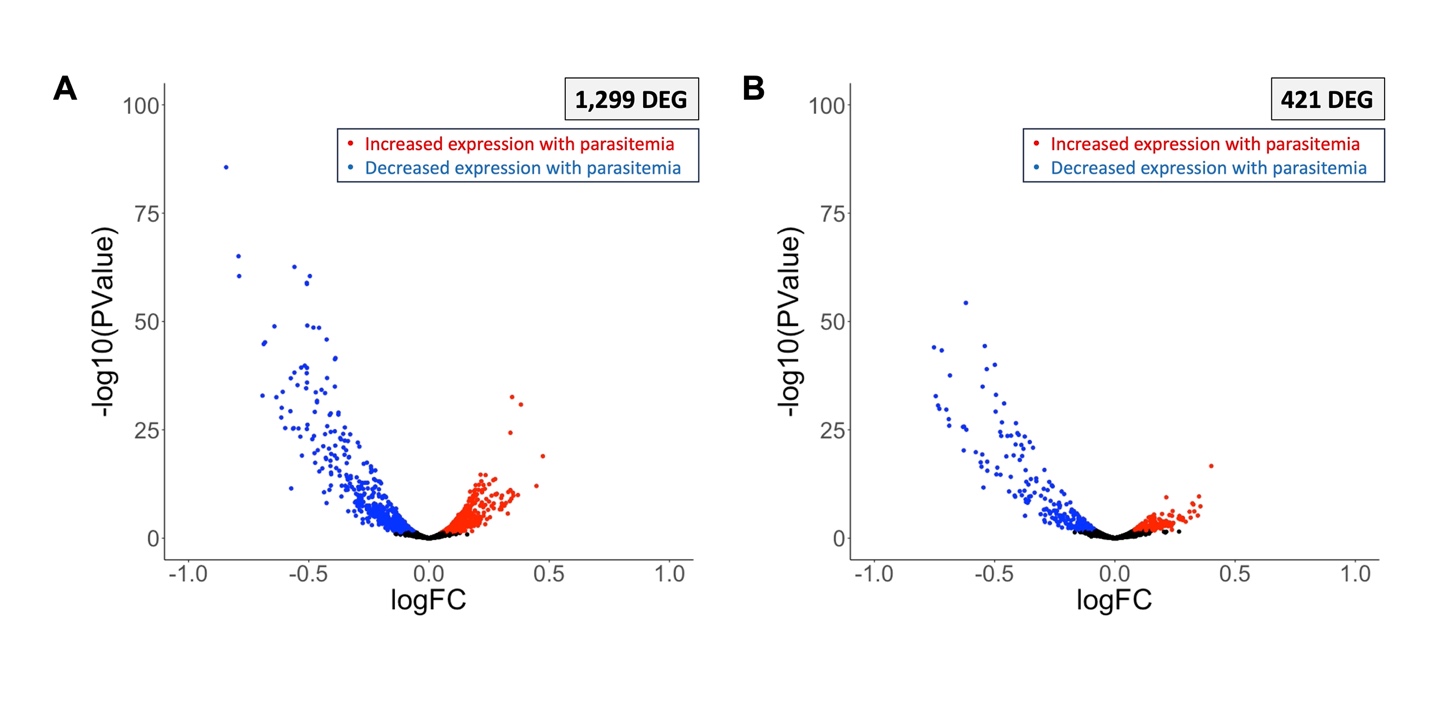

**Supplemental Figure 5: Some parasite genes are correlated with parasitemia, separate from age.** Volcano plot of *P. falciparum* genes associated with log(parasitemia) in four- to five-year-old children. Each point represents one gene, plotted by its P-value (y) and log(fold-change) value (x). Blue points are genes that are significantly negatively associated with log(parasitemia). Red points are genes that are significantly positively associated with log(parasitemia). **A)** Model unadjusted for developmental stage composition; 1,299 significant genes. **B)** Model adjusted for developmental stage composition; 421 significant genes. Significantly differentially expressed genes have an (FDR = 0.1). *DEG = differentially expressed gene.

**Supplemental Figure 6: Human genes that were significantly positively associated with host age at the sequenced infection, mapped onto KEGG Pathways. Green boxes represent genes that are significantly positively and red boxes represent genes that are significantly negatively associated with age at FDR = 0.1. To visualize genes that did change expression in our analysis, but did not meet our significance threshold (FDR = 0.1), light green boxes represent genes that increased in expression with age and light red boxes represent genes that decreased in expression with age at FDR = 0.25. A)** Platelet activation pathway **B)** T cell receptor signaling pathway **C)** B cell receptor signaling pathway **D)** Regulation of the actin cytoskeleton pathway **E)** Focal adhesion pathway.

**Supplemental Figure 7: Human genes that were negatively significantly associated with age at the sequenced infection, mapped onto KEGG Pathways. Green boxes represent genes that are significantly positively- and red boxes represent genes that are significantly negatively associated with age at FDR = 0.1. To visualize genes that did change expression in our analysis, but did not meet our significance threshold (FDR = 0.1), light green boxes represent genes that increased in expression with age and light red boxes represent genes that decreased in expression with age at FDR = 0.25. A)** NOD-like receptor signaling pathway **B)** Toll-like receptor signaling pathway **C)** Antigen processing and presentation pathway **D)** Phagocytosis pathway.

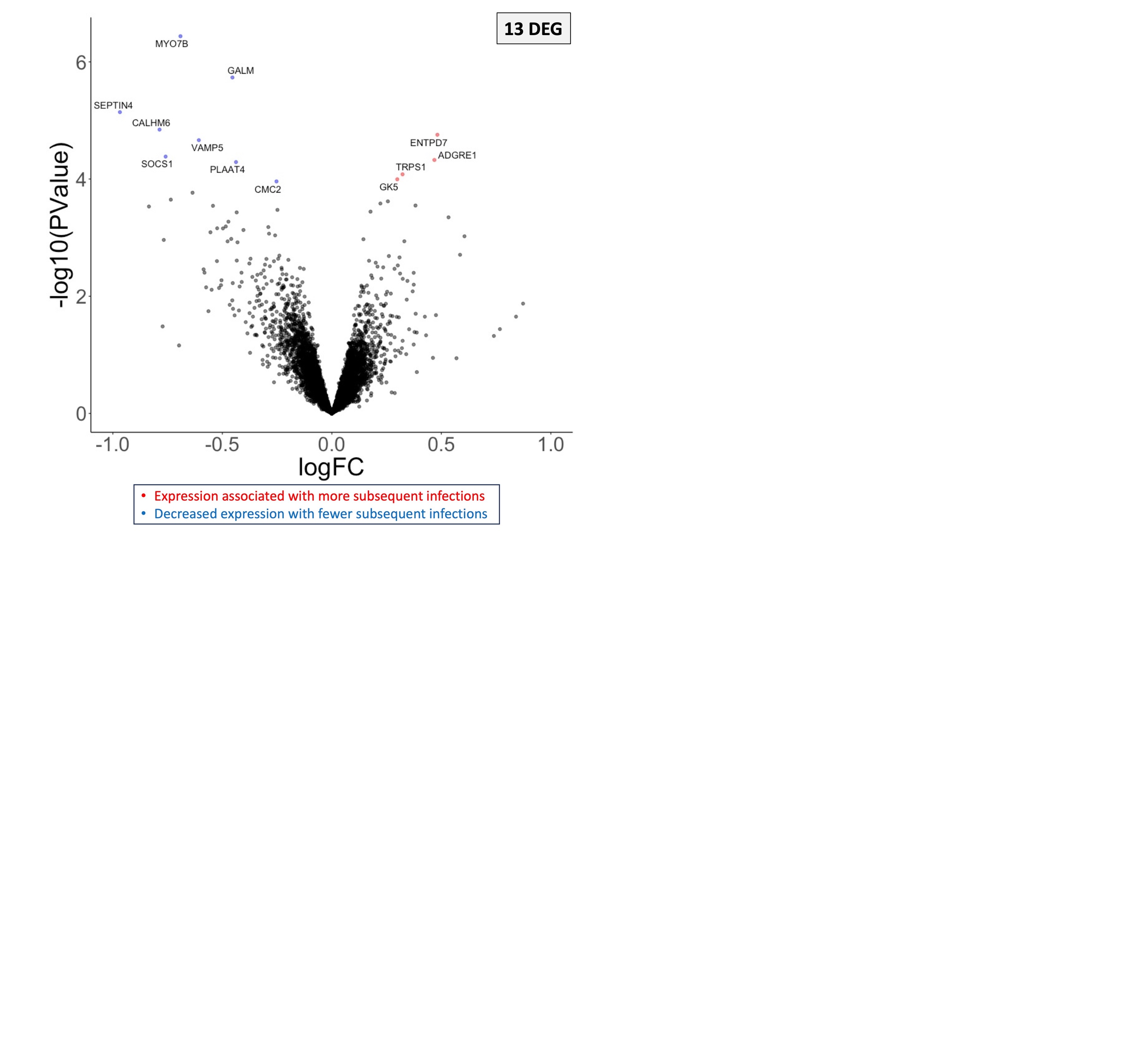

**Supplemental Figure 8: Few human genes are associated with the number of subsequent symptomatic infections in the study period.** Volcano plot of human genes associated with the number of subsequent symptomatic infections in the study period, weighted for the risk of infection during a given time period. Each point represents one gene, plotted by its P-value (y) and log(fold-change) value (x). Blue points are genes that are more highly expressed in children who experienced fewer infections. Red points are genes that are more highly expressed in children who experienced more infections. Significantly differentially expressed genes have an (FDR = 0.1). *DEG = differentially expressed gene.

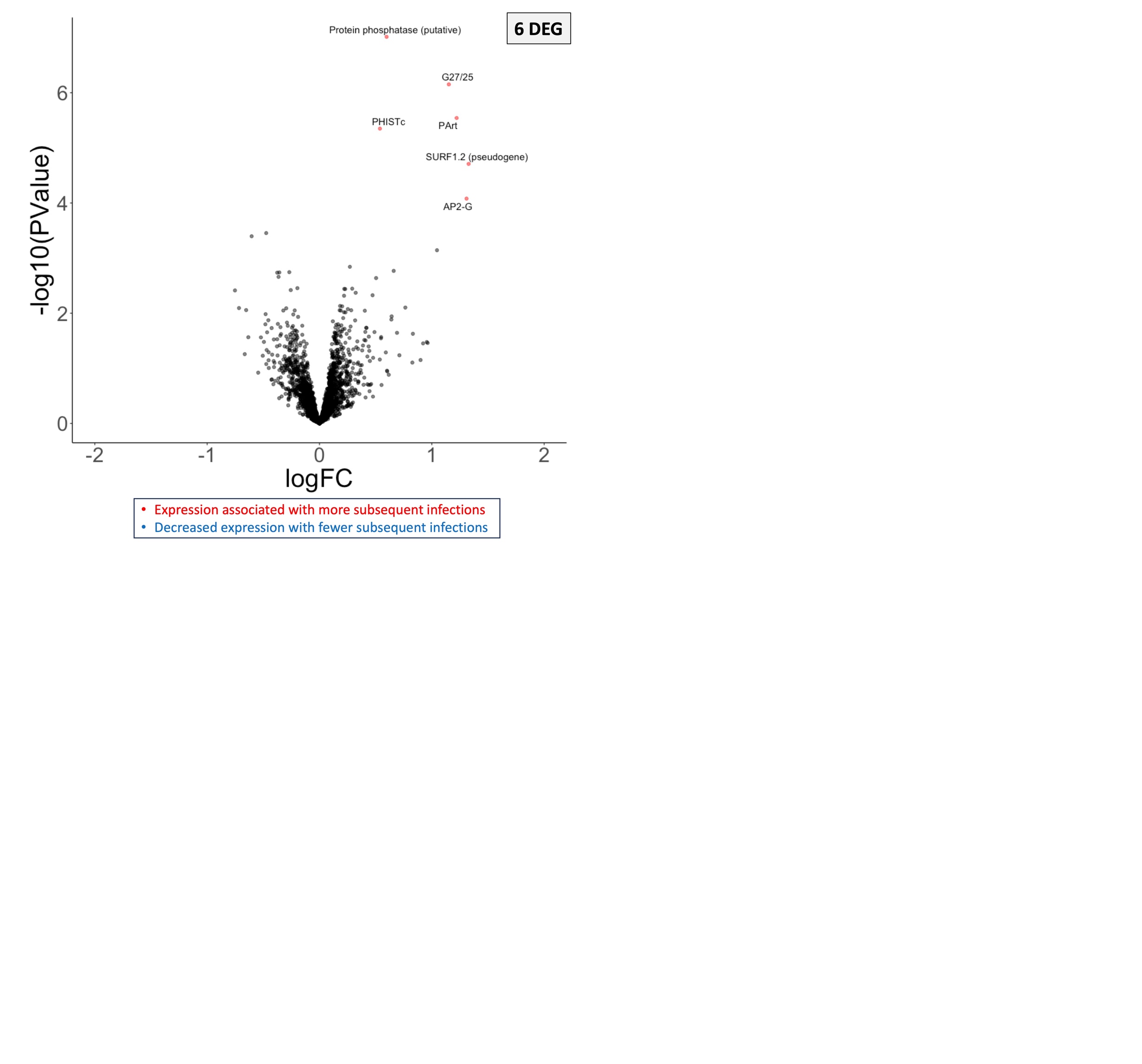

**Supplemental Figure 9: Few parasite genes are associated with the number of subsequent symptomatic infections in the study period.** Volcano plot of parasite genes associated with the number of subsequent symptomatic infections in the study period, weighted for the risk of infection during a given time period. Each point represents one gene, plotted by its P-value (y) and log(fold-change) value (x). Blue points are genes that are more highly expressed in parasites infection children who experienced fewer infections. Red points are genes that are more highly expressed in parasites infection children who experienced more infections. Significantly differentially expressed genes have an (FDR = 0.1). *DEG = differentially expressed gene.
